# From retina to motoneurons: a substrate for visuomotor transformation in salamanders

**DOI:** 10.1101/2022.04.25.489378

**Authors:** Aurélie Flaive, Dimitri Ryczko

## Abstract

The transformation of visual input into motor output is essential to approach a target or avoid a predator. In salamanders, visually-guided orientation behaviors have been extensively studied during prey capture. However, the neural circuitry involved is not resolved. Using salamander brain preparations, calcium imaging and tracing experiments, we describe a neural substrate through which retinal input is transformed into spinal motor output. We found that retina stimulation evoked responses in reticulospinal neurons of the middle reticular nucleus, known to control steering movements in salamanders. Microinjection of glutamatergic antagonists in the optic tectum (superior colliculus in mammals) decreased the reticulospinal responses. Using tracing we found that retina projected to the dorsal layers of the contralateral tectum, where the dendrites of neurons projecting to the middle reticular nucleus were located. In slices, stimulation of the tectal dorsal layers evoked glutamatergic responses in deep tectal neurons retrogradely labelled from the middle reticular nucleus. We then examined how tectum activation translated into spinal motor output. Tectum stimulation evoked motoneuronal responses, which were decreased by microinjections of glutamatergic antagonists in the contralateral middle reticular nucleus. Reticulospinal fibers anterogradely labelled from tracer injection in the middle reticular nucleus were preferentially distributed in proximity with the dendrites of ipsilateral motoneurons. Our work establishes a neural substrate linking visual and motor centers in salamanders. This retino-tecto-reticulo-spinal circuitry is well positioned to control orienting behaviors. Our study bridges the gap between the behavioral studies and the neural mechanisms involved in the transformation of visual input into motor output in salamanders.

## Introduction

Visually-guided orientation behaviors allow animals to move towards or away from a stimulus. In salamanders, visually-guided orientation behaviors have been extensively studied in the context of prey catching. When a prey appears in the visual field, such as a cricket (Schülert and Dicke 2002) or a worm (Roth, 1987), some salamander species turn their head towards the prey and use full body movements to follow it and eventually snap it (Schülert and Dicke 2002). Other salamander species ambush their prey and use motion extrapolation to catch it by projecting their tongue (Borghuis and Leonardo 2015). When presented with two different quantities of prey on the left and right visual fields, salamanders orient towards the highest number of preys (Uller et al. 2003, Krusche et al. 2010). However, the circuitry underlying asymmetric visually-guided orientation behaviors is not fully resolved in salamanders.

A brain region playing a key role in the integration of visual inputs to generate eye, head or body movements to approach or avoid a sensory object is the optic tectum, also called superior colliculus in mammals (for review Isa et al. 2021, Basso et al. 2021). In salamanders as in other vertebrates, the anatomical organisation of the retinal inputs in the tectum follows the topography of the visual field, and therefore builds a map of visual inputs (Grüsser-Cornehls and Himstedt 1973, Margolis 1976ab, Manteuffel et al. 1989, Jakway and Riss 1972, Gruberg 1973, Guillery and Updyke 1976, Caldwell and Bergman 1977, Fritzsch 1980, Fujisawa et al. 1981, Ingham and Güldner 1981, Rettig and Roth 1982, Rettig 1988). Extracellular recordings showed that salamander tectal neurons respond to specific size, contrast or movement pattern of a stimulus moving in the visual field (Grüsser-Cornehls and Himstedt 1973, Roth 1982, Schuekert and Dicke 2005). Single cell fills and immunohistochemistry provided detailed information about the anatomy and overall projection patterns of salamander tectal neurons (Dicke 1999, Roth et al. 1999, Roth et al. 1990, Landwehr and Dicke 2005). However, the precise downstream targets through which tectal neurons generate asymmetrical body movements are largely unknown in salamanders.

In other vertebrates, tectal output neurons project down to command neurons called reticulospinal (**RS**) neurons, which can induce orientation movements through projections to spinal motor circuits (e.g. lamprey: Ullén et al. 1997, Zompa and Dubuc 1996, 1998, Saitoh et al. 2007, Kardamakis et al. 2015, 2016, Pérez-Fernández et al. 2017, Suzuki et al. 2019, Kamali Sarvestani et al. 2013, Kozlov et al. 2014; zebrafish: Roeser and Baier 2003, Gahtan et al. 2005, Bianco and Engert 2015, Helmbrecht et al. 2018, for review Bollmann 2019; mice: Hoy et al. 2019, Cregg et al. 2020, Usseglio et al. 2020, and many other vertebrates for review see Basso et al. 2021). In salamanders, a region that contains many RS neurons and controls asymmetric steering movements is the middle reticular nucleus (**mRN**) (Ryczko et al. 2016a,b,c). In salamander semi-intact preparations, where the brain is accessible and the body moves in the recording chamber, unilateral activation of the mRN evokes ipsilateral bending movements. Increase in stimulation strength increases the bending angle (Ryczko et al. 2016c, for review Ryczko et al. 2020b). Anatomical studies suggest that descending tectal projections target reticular nuclei (Finkenstädt et al. 1983, Naujoks-Manteuffel and Manteuffel 1990), but whether these projections target RS neurons and provide physiological input is not resolved. Two modeling studies successfully generated prey detection and orientation movements during ongoing locomotion in a salamander neuromechanical model controlled by a simulated neural circuit comprising the retina, tectum, RS neurons, and spinal motoneurons (Ijspeert and Arbib 2000, Petreska 2004; see also previous models of Eurich et al. 1992, 1995). However, the existence of this circuitry was not tested experimentally in salamanders.

Here, we aimed at uncovering the neural circuitry through which visual input is transformed into an asymmetric motor command in salamanders. Using tracing and calcium imaging, we found a retino-tecto-reticulo-spinal pathway that constitutes a substrate for visually-guided orientation movements in salamanders. Our study opens the door to mechanistic and computational studies of visuomotor transformations in salamanders. It also contributes to our understanding of visuomotor transformations together with other models such as e.g. lamprey, zebrafish, rodent and primate (Rozenblit and Gollisch 2020). In addition, the regenerative capacities of the salamander make it a unique model to examine how this circuitry is rewired after injury (Joven and Simon, 2018, Rozenblit and Gollisch 2020).

## Materials and Methods

### Ethics statement

The procedures were in accordance with the guidelines of the Canadian Council on Animal Care and approved by the animal care and use committees of the Université de Sherbrooke.

### Animals

We used juvenile salamanders *Pleurodeles waltl* (Andras Simon laboratory, Karolinska Institute, Stockholm, Sweden) of either sex with snout–vent length (SVL) ranging from 50 to 80 mm. The animals were housed in plastic tanks at 17-19°C degrees and fed with bloodworms, beef meat and fish pellets. The anatomical experiments were carried out in 8 CAG:NucCyt females and 1 CAG:NucCyt male and 3 wild-type females. The physiology experiments were carried out in 15 leucystic males and 9 leucystic females and 2 wild-type females.

### Surgical procedures

The animals were anesthetized with tricaine methanesulfonate (MS-222, 200 mg/mL, Sigma) and transferred into an oxygenated Ringer’s solution (in mM 130 NaCl, 2.1 KCl, 2.6 CaCl2, 1.8 MgCl2, 4 HEPES, 5 glucose, and 1 NaHCO3, pH = 7.4) at room temperature. The skin and muscles with removed around the cranium with a fine forceps and spring scissors (Fine Science Tools). To provide access to the retina, the cornea, iris and lens were removed. To provide access to the first 5 spinal segments, a dorsal laminectomy was performed. The preparations were then prepared for either anatomy or physiology experiments.

### Anatomical tracing

To retrogradely label RS neurons, a coronal transection was made on one side at the level of the second spinal segment using a fine microscalpel blade. To retrogradely label tectal neurons, a coronal transection was made on one side at the level of the middle reticular nucleus using a similar microscalpel blade. To anterogradely label the retinal inputs, the optic nerve was sectioned and the tracer was placed on the cut axons. In all cases the gap created by the lesion was filled with Texas Red Dextran Amine or biocytin for 15 minutes allowing the tracer to fill the axons. The injection site was then rinsed with Ringer to remove the excess of tracer and the preparations was pinned down at the bottom of a sylgard chamber continuously perfused with oxygenated Ringer’s solution for 4 hours to allow retrograde transport of the tracer. The preparation was then fixed in 4% paraformaldehyde for 24h at 4°C and transferred in a phosphate buffer solution (0.1M) containing 0.9% of NaCl (PBS, pH = 7.4) containing 4% (wt/vol) of paraformaldehyde (PFA 4%). The preparation was then incubated in a PBS solution containing 20% (wt/vol) sucrose for 24 h.

### Immunofluorescence

The procedures were as previously reported (Ryczko et al. 2013, 2016a, 2016b, 2017, 2020a, Flaive et al. 2020a, Fougère et al. 2021a,2021b, van der Zouwen et al. 2021). The brains were snap frozen in 2-methylbutane (-45°C ± 5°C). All steps were carried out at room temperature unless stated otherwise. Brain sections (40 µm thickness) were obtained at -20°C using a cryostat (Leica CM 1860 UV) and collected in PBS (0.1M, pH 7.4, NaCl 0.9%). The sections were rinsed three times during 10 min in PBS and incubated during 1 h in a blocking solution containing 5% (vol/vol) of normal donkey serum and 0.3% Triton X-100 in PBS. Sections were incubated at 4°C for 48 h on an orbital shaker (40 rpm) in a PBS solution containing the primary antibody against glutamate Islet-1 and Islet-2 homeobox (Islet1/2) [mouse anti-Islet-1/2, Developmental Studies Hybridoma Bank (DSBH) 39.4D5, lot 1ea-24 g/mL (1:100), RRID AB_2314683] (Flaive et al. 2020a). Next, sections were rinsed three times in PBS and incubated for 4 h in a solution containing a secondary antibody to reveal Islet1/2 [donkey anti-mouse Alexa Fluor 647, Invitrogen A-31571 lot 2136787 (1:400), RRID AB_ 162542]. Biocytin was revealed by rinsing the slices three times with for 10 min in PBS and incubating them 30 min in PBS containing streptavidin-Alexa Fuor 488 (Invitrogen S32354, lot 18585036 [1:500]). The sections were rinsed three times in PBS for 10 min and mounted on Colorfrost Plus (Fisher) with a medium without DAPI (Vectashield H-1000), covered with a 1.5 type glass coverslip, and stored at 4°C.

### Antibody specificity

As described elsewhere (Flaive et al. 2020a), the DSBH 39.4D5 supernatant was successfully used to label Islet-1/2, a well-established marker of motoneuron identity (Ericson et al. 1992; Hutchinson and Eisen 2006; Tsuchida et al. 1994). It has been used to label motoneurons in salamanders (Moreno and González 2007; Moreno et al. 2018, Flaive et al. 2020a), zebrafish (Hutchinson and Eisen 2006) chickens, mice, and rats (Yamamoto and Henderson 1999). The antibody detects Islet-1 and Islet-2 (Islet-1/2) proteins, and this is consistent with the patterns of mRNA labeling using in situ hybridization (Hutchinson and Eisen 2006). Western blots showed that this antibody labels the Islet-1 protein proportionally with the level of Islet-1 mRNA quantified with RT-PCR (Liu et al. 2011).

### Microscopy

Whole mounts or brain sections were observed using an epifluorescent microscope (Zeiss AxioImager M2 bundled with StereoInvestigator 2018 software v1.1, MBF Bioscience) or a confocal microscope (Leica TCS SP8 nanoscope bundled with LASX software (Leica). For confocal stacks, we used a 40X objective, 1,024 × 1,024 pixel resolution (290.62 × 290.62 µm), 3 frames averaging, bidirectional scanning at 400-Hz speed and pinhole opened at 65.3 µm. The number of images taken for a stack varied from 146 to 208 and the total depth acquired varied from 24.93 to 35.65 µm. Contrast levels were adjusted so that all fluorophores were visible simultaneously, and digital images were merged using Photoshop CS6 (Adobe).

### Optical density

To estimate the density of the descending RS innervation in the spinal cord, we measured in spinal cord sections the optical density of the fluorescent signal generated by the fibers anterogradely labelled from TRDA injection in the reticular nucleus (Xavier et al. 2005, Perlbarg et al. 2018) as we did in a previous study (Fougère et al. 2021b). For each animal, ten coronal sections at the level of the spinal cord were imaged using an epifluorescent microscope Zeiss AxioImager M2. Using ImageJ, regions of interest were drawn manually over the photographs to delineate the ventral spinal cord. To estimate optical density in the regions of interest, the photographs were converted into a greyscale and compared to a calibrated greyscale taken from the ImageJ optical density calibration protocol (https://imagej.nih.gov/ij/docs/examples/calibration/) (Xavier et al. 2005, Fougère et al. 2021b).

### Tectum slices

To record the tectum cells projecting to the mRN, the tracer Ca^2+^ green was injected at the level of the mRN bilaterally and pinned down in a cold (10°C degrees) oxygenated chamber overnight. The next day, coronal slices were obtained as previously described (Flaive and Ryczko 2020). Briefly, the brain was glued at the level of the transverse section rostral to the optic nerve onto the specimen disk and placed in the slicing chamber of a VT1000S vibrating-blade microtome (also called vibratome, Leica) filled with the ice-cold sucrose-based solution. (in mM: 3 KCl, 1.25 KH2PO4, 4 137 MgSO4, 26 NaHCO3, 10 Dextrose, 0.2 CaCl2, 219 Sucrose, pH 7.3–7.4, 300-320 138 mOsmol/kg) bubbled with 95% O2 and 5% CO2, with the vibratome blade facing the dorsal side of the brain. Coronal slices (350 µm thickness) were prepared using high frequency blade oscillations (100 Hz, scale setting “10”) and slow blade progression (0.15 mm/s, scale setting “3”), under visual inspection with a stereomicroscope (Leica) installed over the VT1000S. To lessen the brain movements evoked by blade vibrations, a small brush was gently positioned against the ventral side of the brain, at the level of the slice being cut. Slices were then allowed to rest at room temperature for an hour in a chamber filled with aCSF (in mM: 124 NaCl, 3 KCl, 1.25 KH2PO4, 1.3 MgSO4, 26 NaHCO3, 10 Dextrose, and 1.2 CaCl2, pH 7.3–7.4, 290–300 mOsmol/kg) bubbled with 95% O2. Brain slices were carefully placed at the bottom of the recording chamber with the brush under the microscope and secured in place with one or two platinum wires disposed on the slice.

### Calcium imaging

The procedure was similar to that we previously used in lampreys and salamanders (e.g. Brocard et al. 2010; Ryczko et al. 2016a,b,c). The brain rostral to the optic nerve was removed following a transverse section. The RS neurons were retrogradely labeled by placing crystals of Ca^2+^ Green dextran amine (MW,3,000 Da, Invitrogen) at the level of the first segment of the spinal cord after a transection to record RS neurons in some experiments, or at the level of the cervical segment (third ventral root) to record motoneurons in other experiments. The preparation was transferred to a cold (10°C degrees) and oxygenated chamber for 18–24 h to allow the dye to retrogradely labelled neurons. The following day, the preparation was pinned down dorsal side up to the bottom of a recording chamber covered with Sylgard (Dow Corning, Midland, MI) and perfused with oxygenated Ringer’s solution (1.67 mL/min) at room temperature. A stimulation electrode was placed on one side in the retina or in the optic tectum. The RS nuclei were easily identified under the microscope on the basis of our previous reports describing the distribution of RS cells in salamanders (Naujoks-Manteuffel and Manteuffel, 1988; Sanchez-Camacho et al., 2001, Ryczko et al. 2016a,b,c). An optimal focal plane was chosen for imaging cells in the mRN. For motoneuron recordings, the spinal cord was delicately opened for the dorsal side with spring scissors, and the dorsal part of the spinal cord was removed ipsilaterally to the labelled motoneurons. The cervical motoneurons were easily identified from the ventral root injection. To measure the change in fluorescence in neurons, regions of interest were manually delineated around the cell bodies labeled with the Ca^2+^ dye using the Region of Interest Manager in ImageJ. Ca^2+^ responses of RS neurons to retina or tectal stimulation were acquired at the rate of two images per second (exposure time of 36 ms) with an Axio Examiner Z1 epifluorescent microscope (Zeiss, Toronto, ON, Canada) coupled with a Colibri 7 illumination system and an ORCA-Flash 4.0 Digital CMOS Camera V3 (Hamamatsu Photonics, Hamamatsu, Japan), with 475 nm laser power set to 8%. Signal analyses were carried out using Excel, Sigmaplot and Matlab. To eliminate the Ca^2+^ signal drift, we subtracted an exponential equation fitted on the fluorescence signal recorded in each cell using Sigmaplot (Nanou et al. 2013). The Ca^2+^ responses were expressed as the relative change in fluorescence (dF/F). For each cell, baseline was defined as the averaged fluorescence before stimulation. All quantifications of RS responses were done using the dF/F peak during the evoked response. The colour plots used to illustrate the variations of fluorescence for each cell as a function of time were generated using the imagesc function in Matlab (Mathworks).

### Drugs

Chemicals were purchased from Sigma, Tocris Bioscience or Invitrogen. In some experiments, the following drugs were bath applied: the AMPA/kaïnate receptor antagonist 6-cyano-7-nitroquinoxaline-2,3-dione (CNQX, 0.9 μM), the NMDA receptor antagonist (2R)-amino-5-phosphonovaleric acid (AP5, 3.5 μM). In some experiments, these antagonists were applied locally in the tectum or in the mRN with glass micropipettes (tip diameter around 3-20 μm) using 30 to 90 pressure pulses (5 psi, 30 ms duration) applied with a Picospritzer (Parker). The injected volumes were estimated by measuring the diameter of a droplet ejected in air from the tip of the pipette multiplied by the number of pressure pulses, and the resulting number of moles ejected was calculated (Le Ray et al., 2003; Ryczko et al., 2013, 2016a,b, 2017, 2020a, 2021).

### Electrical stimulation

Glass-coated tungsten microelectrodes (0.7–3.1 MΩ with 10–40 μm exposed tip) and a Grass S88 stimulator coupled to a Grass PSIU6 photoelectric isolation unit for controlling stimulation intensity (Astro Med) were used. The stimulation electrode was placed in the retina or optic tectum. In tectal slices, the electrode was placed in the dorsal layers, ∼100 μm lateral to the level of the recording site of the deeper located tectal output neurons, to elicit action potentials in retinofugal axons contacting the recorded tectal output neurons. The electrical stimulation consisted of square pulses (2 ms duration) applied with a frequency of 10 Hz for 5 s. A pause of at least 2 min was given between two stimulation trains. The stimulation intensities ranged from 1 to 70 μA.

### Statistical analysis

Data are presented as mean ± standard error of the mean (**SEM**) unless stated otherwise. Statistical analyses were done using Sigma Plot 12.1. Parametric analyses were used when assumptions for normality and equal variance were respected, otherwise non-parametric analyses were used. To compare the means between two dependent groups, a two-tailed paired t-test or a nonparametric Wilcoxon signed rank test was used. To compare the means between two independent groups, a two-tailed t-test or a nonparametric Mann– Whitney rank sum test was used. Linear and regressions between variables, their significance, and the confidence intervals were calculated using Sigma Plot 12.0. Statistical differences were assumed to be significant when *P* < 0.05.

## Results

### Retina stimulation evokes RS responses

In salamanders, RS neurons of the middle reticular nucleus (**mRN**) evoke ipsilateral steering movements when activated pharmacologically in semi-intact preparations (Ryczko et al. 2016c). We examined whether retina could evoke responses in mRN RS neurons retrogradely labelled by injection of the calcium (Ca^2+^) sensor Ca^2+^ green in the first spinal segment (Ryczko et al. 2016a,b,c). An electrical stimulation electrode was placed in the retina on one side in the isolated brain (Fig. 1A). Trains of electrical stimulation applied to the retina evoked Ca^2+^ responses in mRN RS neurons. The RS responses were stronger ipsilateral to the stimulated retina in single animal data (Fig. 1B-F, G,H). We pooled the data from 6 preparations and normalized in each preparation the Ca^2+^ responses as a percentage of the maximal response recorded, and the stimulation intensity as a percentage of the maximal stimulation intensity used (e.g. Ryczko et al. 2016a, 2017, van der Zouwen et al. 2021). In the pooled data as well, the responses evoked by retina stimulation were stronger in RS neurons ipsilateral to the stimulated retina (Fig. 1I-J). At the highest stimulation intensity for each animal, the responses on the ipsilateral side were 31% stronger than on the contralateral side (53.8 ± 7.5% ipsi vs. 22.4 ± 3.9% of the maximal response, n = 49 ipsilateral RS cells vs. n = 41 contralateral RS cells pooled from n = 6 preparations). On both sides, we found a linear positive relationship between the retina stimulation intensity and the peak of RS Ca^2+^ responses (linear fit, *P* < 0.001, Fig. 1K-L data pooled from 6 preparations). This indicates that a neural circuitry transformed retina activation on one side into an ipsilaterally stronger RS descending command.

**Figure 1.**
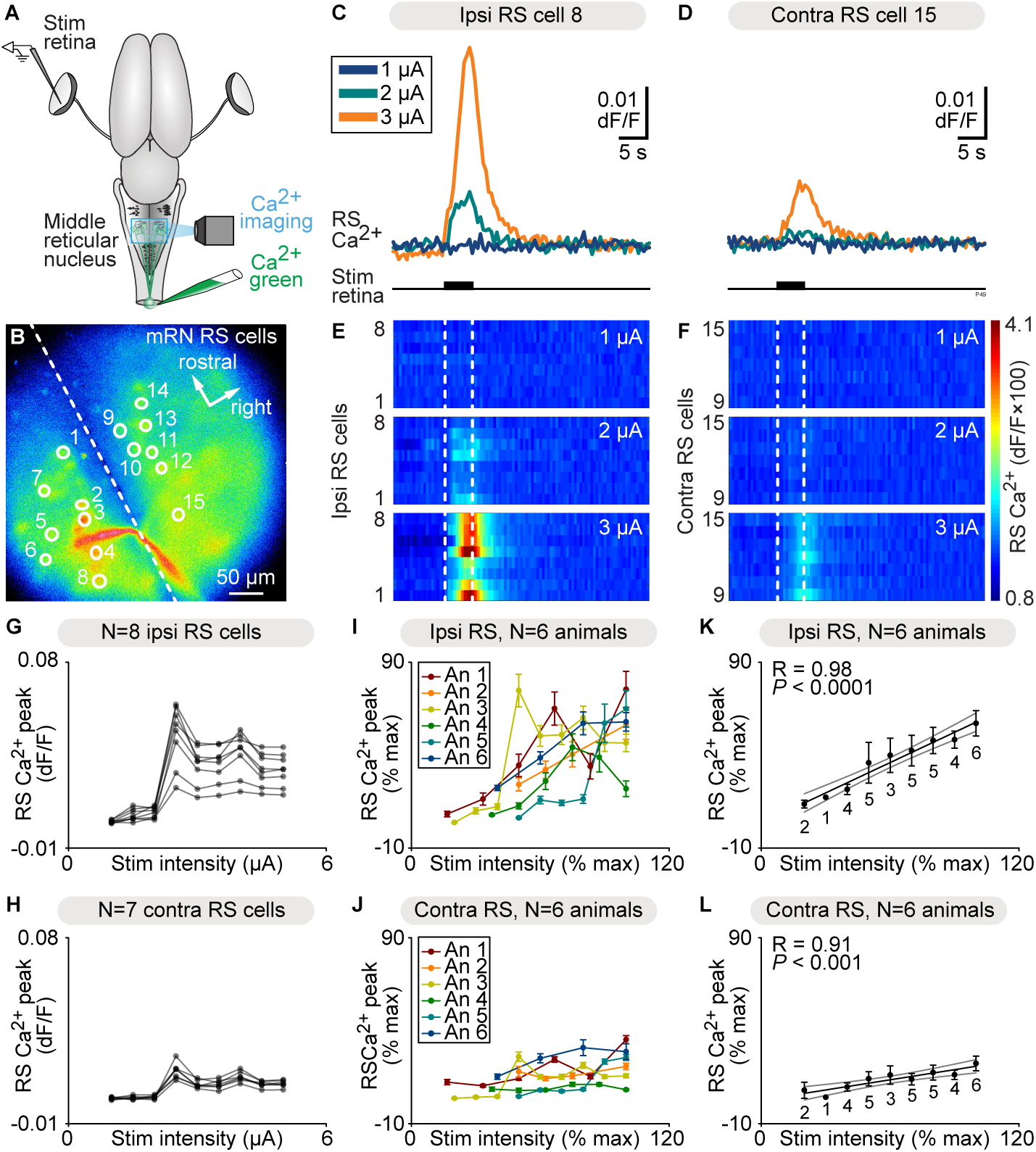
Retina stimulation evokes stronger reticulospinal (RS) responses on the ipsilateral side. **A.** Schematic dorsal view of a salamander brain. A calcium (Ca^2+^) sensor (calcium green) was applied onto the first spinal segments to retrogradely label RS neurons on both sides. A stimulation electrode was placed in the retina on one side and optical recordings of the RS neurons of the middle reticular nucleus (mRN) were obtained. **B.** Ca^2+^ fluorescence at rest of mRN RS neurons. **C-D.** Representative Ca^2+^ response of individual RS cells on the ipsilateral side (C) and contralateral side (D) of the mRN in response to retina stimulation on left side (1-3 µA, 5 s train, 10 Hz, 2 ms pulses). **D-F.** Color plots showing the Ca^2+^ responses of mRN RS cells for increasing retina stimulation intensities. Each line illustrates the response of one cell, with cells 1 to 8 located ipsilateral to retina stimulation (D) and 9 to 15 located contralateral to retina stimulation (F). Onset and offset of stimulation are indicated with vertical white dotted lines. Warm colors (red) indicate stronger Ca^2+^ responses. **G-H.** Relationships between Ca^2+^ response peak and retina stimulation intensity in RS cells illustrated in B-F. Each trace represents the responses of a single RS cell ipsilateral to stimulation (G) or contralateral to stimulation (H). **I-J.** Relationships in 6 preparations between Ca^2+^ response peak (mean ± SEM) and retina stimulation intensity for RS cells located ipsilateral (n = 49 RS cells, 4 to 11 cells per preparation, I) and contralateral (n = 41 RS cells, 4 to 11 cells per preparation, J) to the stimulated retina (0.6-30 µA, 5 s train, 10 Hz, 2 ms pulses). In each preparation, response peaks were expressed in % of the maximal peak recorded, and stimulation intensity in % of the maximal intensity used. **K-L.** Relationship between Ca^2+^ response peak (mean ± SEM) and stimulation intensity in the same 6 animals as in I-J. Data was binned with a bin size of 10%. The data followed a linear polynomial function both for ipsilateral RS cells (solid black line, *P* < 0.001, R = 0.98, K) and contralateral RS cells (solid black line, *P* < 0.001, R = 0.91, L). Note that RS cells contralateral to retina stimulation generated lower maximal responses. Grey lines illustrate the confidence intervals.

### Retina projects to thalamus, pretectum and tectum

We then examined the circuitry involved in this sensorimotor transformation. To identify the central targets of the retina, we anterogradely labelled the retinofugal projections by injecting a tracer in the optic nerve (biocytin, n = 2 animals; TRDA, n = 1 animal) (Fig. 2A). These injections anterogradely labelled the optic tract (Fig. 2B), and fibers in the contralateral lateral geniculate nucleus of the thalamus (*corpus geniculatum thalamicum*) and in the neuropil of Bellonci (Fig. 2C). More caudally in the brain, anterogradely labelled fibers were found in the contralateral pretectum (Fig. 2D), and a dense innervation was found in the dorsal layers of the contralateral tectum (Fig. 2E). Occasionally some fibers were observed in the tectum ipsilateral to the injected optic nerve. Some retrogradely labelled cell bodies projecting to the retina were also observed in the tegmentum ventral to the tectum, as reported in many vertebrate species (e.g. ventral brain in Fig. 2E, for review see Repérant et al. 2006, 2007). Altogether, this indicates that the retina sends projections to the thalamus, pretectum and tectum, and receives projections from central neurons in the tectum, features that are well conserved from basal vertebrates to mammals (see discussion, Suryanarayana et al. 2020, Repérant et al. 2006, 2007).

**Figure 2.**
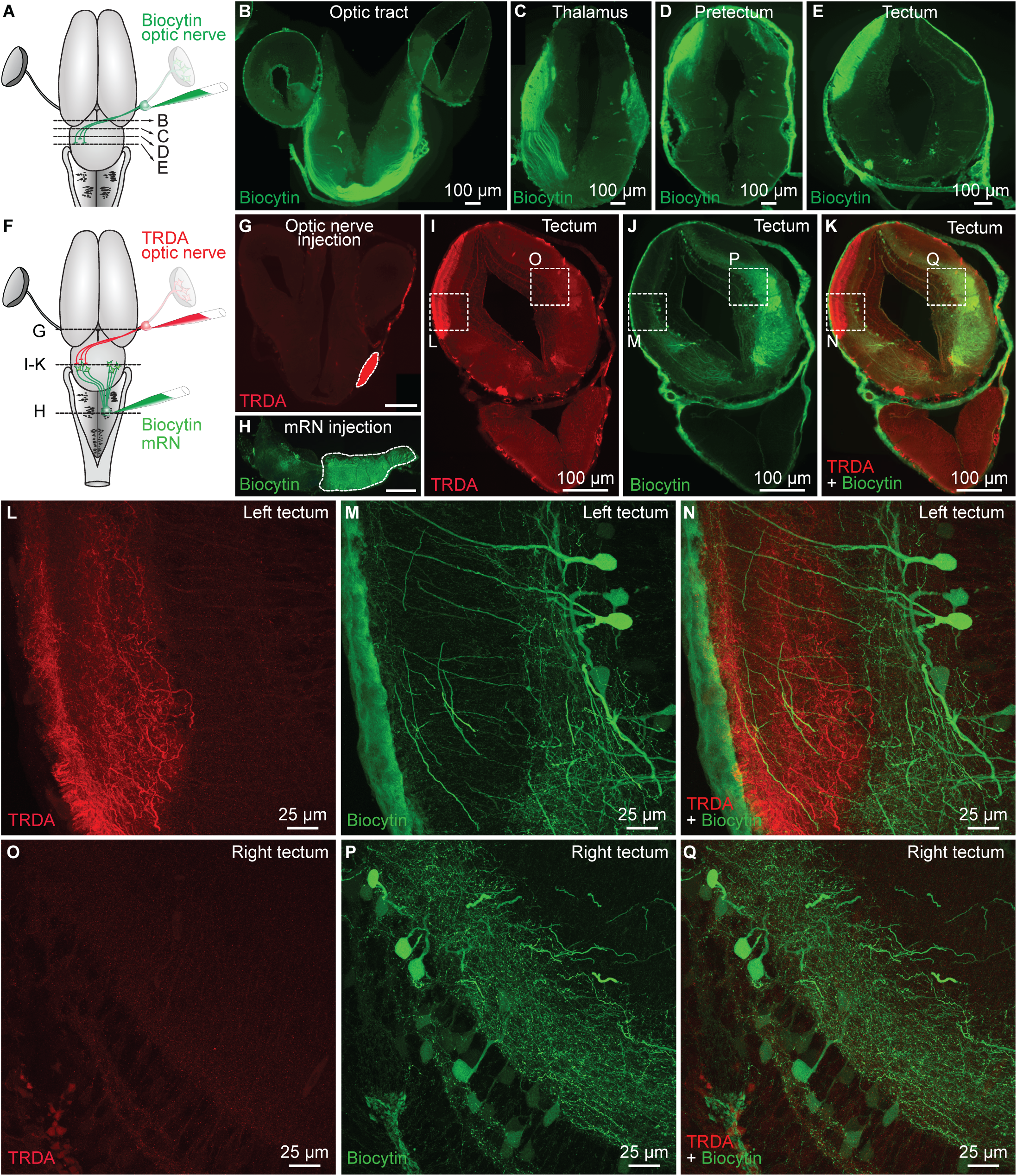
Retinofugal fibers innervate the dendrites of tectal neurons projecting down to the middle reticular nucleus. **A.** Scheme showing a dorsal view of the salamander brain. A tracer (biocytin, green) was injected in the right optic nerve. After 4h of tracing, the brain was fixed and coronal sections were obtained. **B-C.** Coronal sections showing that tracer injection in the optic nerve labelled fibers in the optic tract (B), thalamus (C), pretectum (D) and tectum (E). **F-H.** Scheme showing a dorsal view of the salamander brain (F). A first tracer was injected in the right optic nerve (Texas Red Dextran Amine, TRDA, red, G) and a second tracer (biocytin, green, H) was injected on the right side of the middle reticular nucleus (mRN), where many reticulospinal (RS) neurons are located. **I-K.** Coronal sections showing the anterogradely labeled retinofugal projections (TRDA, red) and the retrogradely labelled tectal neurons (biocytin, green). Scale bar in G-H: 100 µm. **L-N.** Confocal magnification of the squares in I-K, showing the presence of retinofugal projections (TRDA, red) in the dorsal layers of the left tectum (i.e. contralateral to the injected optic nerve). In green, neurons in the left tectum retrogradely labelled by biocytin injection in the right mRN. **O-Q.** Magnification of the squares in I-K, showing the rarity of retinofugal projections in the dorsal layers of the right tectum, i.e. ipsilateral to the injected optic nerve. In green, many neurons of the right tectum were retrogradely labelled by biocytin injection in the right mRN.

### Tectal neurons innervated by retinal fibers project to the mRN

The tectum plays a key role in visuomotor transformation in vertebrates (Basso et al. 2021, Isa et al. 2021). Next, we examined using double labelling experiments whether relay neurons in the tectum receive projections from the contralateral retina, and project to mRN RS neurons ipsilateral to the stimulated retina, which show the strongest responses to retina stimulation (Fig. 1). We injected a first tracer in the right retina to anterogradely label retinofugal projections, and a second tracer in the right mRN to retrogradely label tectal output neurons (Fig. 2F-H). In coronal slices of the tectum, we found that retinofugal fibers and varicosities were densely labelled in the tectum contralateral to the retina. There, retinofugal fibers were in proximity with the dendrites of deep tectal output neurons projecting to the contralateral mRN, therefore ipsilateral to the injected retina (n = 3 preparations, Fig. 2I-N). In addition, many tectal neurons were retrogradely labelled ipsilaterally to the mRN injection site, but only rare fibers from the ipsilateral retina were found in their dendritic trees (n = 3 preparations, Fig. 2I-K,O-Q). Altogether this indicated that the retina projects mainly to the contralateral tectum, where tectal output neurons project to the ipsilateral and contralateral sides of the mRN, thus providing a substrate for the bilateral RS responses evoked by unilateral stimulation of the retina.

### Tectal neurons projecting to mRN receive glutamatergic inputs from retina

Next, we examined the transmission involved. In the isolated brain, we stimulated the retina and recorded the Ca^2+^ RS responses in the mRN (Fig. 3A). In each preparation, the Ca^2+^ responses were normalized as a function of the maximal response. Responses were stronger (+37%) on the ipsilateral side (*P* < 0.001, t-test Fig. 3B-C), consistent with the results presented in Fig. 1. The retina-evoked Ca^2+^ responses were largely decreased after bath application of the glutamatergic antagonists CNQX and AP5 in mRN RS located ipsilaterally (*P* < 0.001, paired t test, n = 30 RS cells) and contralaterally (*P* < 0.001, paired t test, n = 30 RS cells) to the stimulated retina (Fig. 3B-C, data pooled from 4 preparations, 5-9 trials per drug condition per preparation).

**Figure 3.**
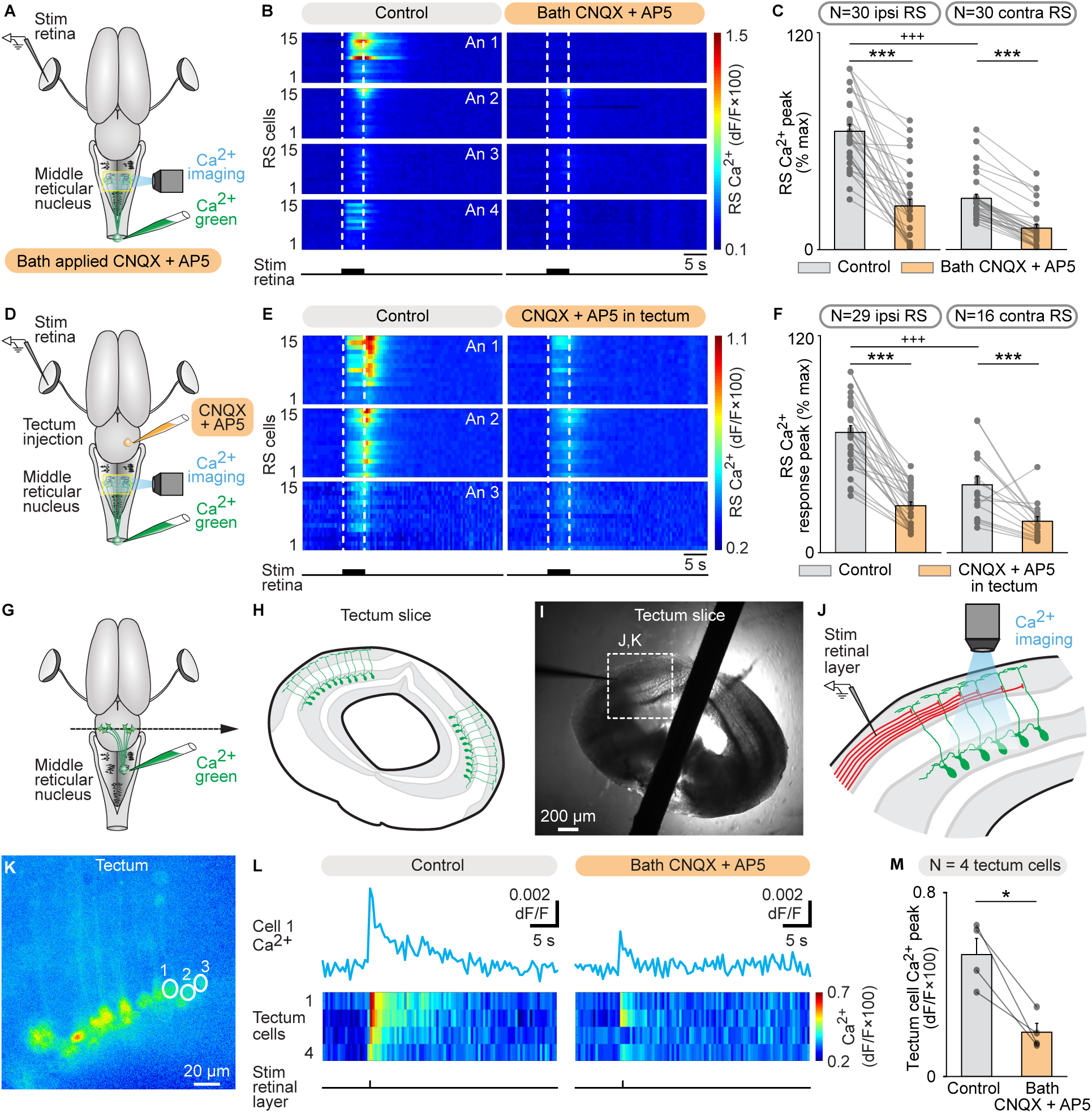
Retina stimulation evokes reticulospinal (RS) responses in the middle reticular nucleus (mRN) via a glutamatergic relay in the contralateral tectum. A. Schematic dorsal view of a salamander brain. A calcium (Ca^2+^) sensor (Ca^2+^ green) was applied onto the first spinal segments to retrogradely label RS neurons. A stimulation electrode was placed in the retina on one side and optical recordings of RS neurons of the middle reticular nucleus (mRN) were obtained. B. Color plots showing the Ca^2+^ responses of mRN RS cells ipsilateral (N = 30 RS cells pooled from 4 animals, 4-9 cells per animal, An) or contralateral (N = 30 RS cells pooled from 4 animals, 6-11 cells per animal) to retina stimulation (5 s train, 10 Hz, 2 ms pulses, 4-70 µA) before and 8-24 min after bath application of CNQX (1 µM) and AP5 (3 µM). Each line illustrates the average response of one cell (5-8 trials per cell per drug condition). Onset and offset of stimulation are indicated with vertical white dotted lines. Warm colors (red) indicate stronger Ca^2+^ responses. C. Bar charts showing the Ca^2+^ responses before and after bath application of the glutamatergic antagonists on RS cells located ipsilateral (ipsi) and contralateral (contra) to retina stimulation. D-F. Similar representation as in A-C, showing the responses of RS cells ipsilateral (N = 29 RS cells pooled from 3 animals, 7-11 cells per animal) or contralateral (N=16 RS cells cells pooled from 3 animals, 4-8 cells per animal) to retina stimulation (5 s train, 10 Hz, 2 ms pulses, 3-10 µA) before and 4-20 min after local injection of CNQX (0.07 pmol) and AP5 (0.03 pmol) in the tectum contralateral to the stimulated retina using a picospritzer (CNQX 1 mM, AP5 0.5 mM). Each line illustrates the average response of one cell (5-7 trials per cell per drug condition). G-H. To record tectal neurons projecting to the mRN, Ca^2+^ green was injected in the mRN and coronal slices of the tectum were obtained. I-J. A stimulation electrode was placed laterally (∼100 µm) from the apical dendrites of the labelled tectal neurons, and the Ca^2+^ responses of tectal neurons were recorded. K. Ca^2+^ fluorescence at rest of tectal neurons. L. Ca^2+^ response in tectal cells in response to stimulation of the retinal input layer before (single 2 ms pulses, 60-70 µA, 5-7 trials per cell per drug condition) and 6-38 min after bath application of CNQX (1 µM) and AP5 (3 µM). On the color plot, each line illustrates the average response of one cell (5-7 trials per cell per drug condition). M. Bar charts showing that decreased Ca^2+^ response of tectal neurons in presence of glutamatergic antagonists (n = 4 tectal cells pooled from 2 slices from 2 animals). *, *P* < 0.05, ***, *P* < 0.001, paired t test; +++ *P* < 0.001, t test.

Next, we examined whether local blockade of a tectal glutamatergic synaptic relay contralateral to the stimulated retina was sufficient to abolish the retina-evoked RS responses. In the isolated brain, we stimulated the retina and recorded the Ca^2+^ responses in mRN RS neurons (Fig. 3D). These responses were again stronger ipsilaterally (+29%), consistent with Fig.1 and Fig. 2A-C. Local microinjections of the glutamatergic antagonists CNQX and AP5 in the tectum located contralateral to the stimulated retina largely decreased Ca^2+^ RS responses ipsilaterally to the stimulated retina (n = 29 RS cells pooled from 3 preparations) and contralaterally to the stimulated retina (n = 16 RS cells) (Fig. 3F). We further tested the presence of a local glutamatergic synapse between retinofugal inputs and tectal output neurons projecting to the RS in another series of experiments. We retrogradely labelled the tectal output neurons projecting to the mRN using Ca^2+^ green injections in the right mRN (Fig. 3G). Coronal slices of the tectum were obtained (Fig. 3H-I) and the retrogradely labelled tectal neurons and their dendritic trees could be observed (Fig. 3K). A stimulation electrode was placed in the dorsal layers of the tectum, where retinal inputs are located, slightly lateral (∼ 100 µm) to the apical dendritic tuft of the recorded tectal output neurons (Fig. 3I-J). Single pulse stimulations of the dorsal layer evoked Ca^2+^ responses in tectal output neurons projecting to the mRN (Fig. 3L) (n = 4 tectal neurons pooled from 2 slices from 2 animals). These responses were decreased by bath application of glutamatergic antagonists (*P* < 0.05, paired t test, Fig. 3M). Altogether these experiments indicated that retinofugal inputs were relayed by a glutamatergic synapse to tectal output neurons projecting to mRN RS neurons.

### Tectum sends bilateral inputs to mRN RS neurons

We then examined whether the tectum could control the activity of mRN RS neurons through such tecto-reticular pathway. In the isolated brain, a stimulation electrode was placed in the tectum on one side and we used Ca^2+^ imaging to record mRN RS neurons retrogradely labeled by Ca^2+^ green injection in the spinal cord (Fig. 4A-B). Trains of electrical stimulation applied to the left tectum evoked Ca^2+^ responses in RS neurons in single animal data (Fig. 4B-F). Interestingly, this time Ca^2+^ responses were stronger contralaterally to the stimulated tectum, in contrast with the stronger responses recorded ipsilaterally to the stimulated retina (Fig. 1). Increasing the stimulation intensity in the tectum increased the response of RS neurons in single animal data (Fig. 4C-H). In data pooled from 4 animals, we found a linear positive relationship between the stimulation intensity and the amplitude of RS responses on the ipsilateral side (R = 0.87, linear fit, *P* < 0.01, n = 28 RS cells, Fig. 4K), and between the stimulation intensity and the amplitude of RS responses on the contralateral side (R = 0.72, linear fit, *P* < 0.05, n = 32 RS cells, Fig. 4L). At the highest stimulation intensity for each animal, the responses were 28% stronger on the contralateral side (*P* < 0.01, t-test, 34.6 ± 8.8% ipsi vs. 62.7 ± 2.1% contra, n = 28 ipsilateral vs. n = 32 contralateral RS cells pooled from n = 4 preparations). The RS responses evoked by tectum stimulation were decreased by bath application of the glutamatergic antagonists CNQX and AP5 (n = 60 RS cells pooled from 4 preparations, Fig. 4M-N). Altogether this indicated that activation of the tectum induced stronger responses in contralateral RS neurons, and that glutamatergic transmission was involved.

**Figure 4.**
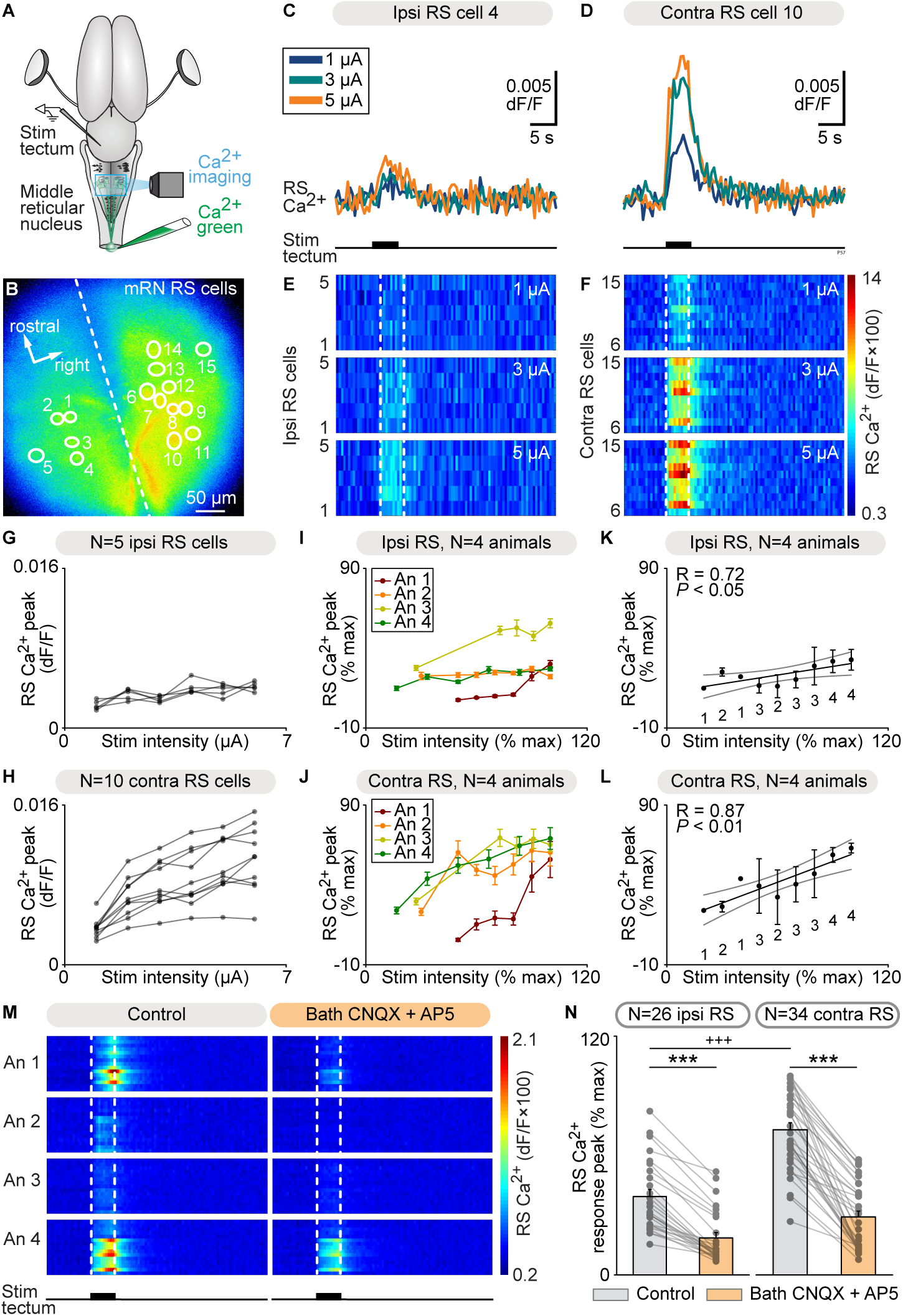
Tectum stimulation evokes stronger reticulospinal (RS) responses on the contralateral side. **A.** Schematic dorsal view of a salamander brain. A calcium (Ca^2+^) sensor (Ca^2+^ green) was applied onto the first spinal segments to retrogradely label RS neurons. A stimulation electrode was placed in the tectum on one side and optical recordings of the RS neurons of the middle reticular nucleus (mRN) were obtained. **B.** Ca^2+^ fluorescence at rest of mRN RS neurons. **C-D.** Ca^2+^ response in individual RS cells on the ipsilateral side (C) and contralateral side (D) of the mRN in response to tectum stimulation on left side (1-5 µA, 5 s train, 10 Hz, 2 ms pulses). **E-F.** Color plots showing the Ca^2+^ responses of mRN RS cells for increasing tectum stimulation intensities. Each line illustrates the response of one RS cell, with cells 1 to 5 located ipsilateral to tectum stimulation (E) and cells 6 to 15 located contralateral to stimulation (F). Onset and offset of stimulation are indicated with vertical white dotted lines. Warm colors (red) indicate stronger Ca^2+^ responses. **G-H.** Relationships between Ca^2+^ response peak and retina stimulation intensity in RS cells illustrated in B-F. Each trace represents the responses of a single RS cell ipsilateral to tectum stimulation (G) or contralateral to stimulation (H). **I-J.** Relationships in 4 preparations between Ca^2+^ response peak (mean ± SEM) and tectum stimulation intensity for RS cells located ipsilateral (n = 28 RS cells, 5 to 8 cells per preparation, I) and contralateral (n = 32 RS cells, 7 to 10 cells per preparation, J) to the stimulated tectum (1-11 µA, 5 s train, 10 Hz, 2 ms pulses). In each preparation, response peaks were expressed in % of the maximal peak recorded, and stimulation intensity in % of the maximal intensity used. **K-L.** Relationship between Ca^2+^ response peak (mean ± SEM) and stimulation intensity in the same 4 animals as in I-J. Data was binned with a bin size of 10%. The data followed a linear polynomial function both for ipsilateral RS cells (solid black line, *P* < 0.05, R = 0.72, K) and contralateral RS cells (solid black line, *P* < 0.01, R = 0.87, L). Note that RS cells contralateral to tectum stimulation generated lower maximal responses. Grey lines illustrate the confidence intervals. **M.** Color plots showing the Ca^2+^ responses of mRN RS cells ipsilateral (N = 26 RS cells pooled from 4 animals, 5-8 cells per animal, An) or contralateral (N = 34 RS cells pooled from 4 animals, 7-10 cells per animal) to tectum stimulation (5 s train, 10 Hz, 2 ms pulses, 1-11 µA) before and 4-20 min after bath application of CNQX (1 µM) and AP5 (3 µM). Each line illustrates the average response of one cell (5-7 trials per cell per drug condition). Onset and offset of stimulation are indicated with vertical white dotted lines. Warm colors (red) indicate stronger Ca^2+^ responses. **N.** Bar charts showing the Ca^2+^ responses before and after bath application of the glutamatergic antagonists on RS cells located ipsilateral (ipsi) and contralateral (contra) to retina stimulation. ***, *P* < 0.001 (paired t test), +++ *P* < 0.001 (t test).

### The mRN projects to spinal motoneurons

We then looked for an anatomical substrate through which the mRN, which receives innervation of the tectum and therefore relays retinal inputs, could feed such inputs to spinal motor circuits to generate body movements. To do this we used dual tracing experiments. To retrogradely label motoneurons innervating axial and forelimb muscles on one side, a tracer (biocytin) was injected in the right ventral root of the cervical segment 3 (Fig. 5A). These motoneurons were positive for the motoneuronal marker Islet- 1/2 (n = 3 preparations, Fig. 5B-E). To label the descending RS projections, a second tracer was injected in the right mRN (Fig. 5F). Such injections anterogradely labeled fibers mostly ipsilaterally in the spinal cord (n = 3 preparations, Fig. 5G-I). We quantified such this asymmetry and found that the optical density of TRDA-labelled fibers was higher ipsilaterally then contralaterally to the mRN injection site (41.9 ± 1.0 ipsilaterally vs. 35.5 ± 1.3 contralaterally, *P* < 0.001, t test, n = 30 sections pooled from 3 preparations, 10 sections per preparation, Fig. 5K). Fibers and varicosities were in apposition with the dendrites of motoneurons (Fig. 5J), suggesting that descending RS projections from the mRN could influence motoneuronal activity.

**Figure 5.**
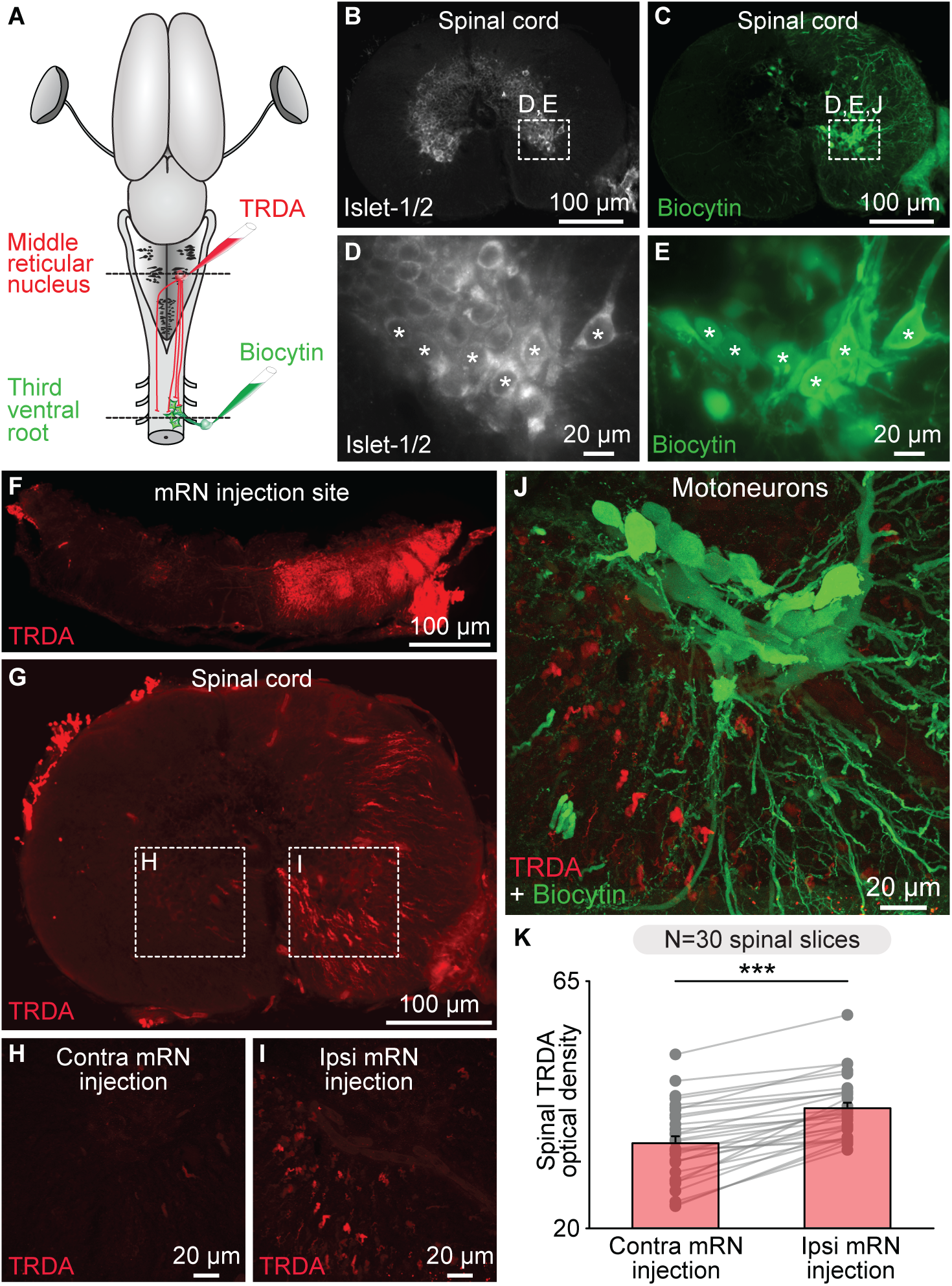
Reticulospinal (RS) neurons of the middle reticular nucleus (mRN) send descending projections to spinal motoneurons on the same side. A-C. Schematic dorsal view of a salamander brain. To anterogradely label the descending RS projections of the mRN, a first tracer (Texas Red Dextran Amine, TRDA, red) was injected in the right side of the mRN. To retrogradely label the motoneurons, a second tracer (biocytin, green) was injected in the right ventral root of the third cervical spinal segment. B-E. Coronal slices of the spinal cord showing that motoneurons retrogradely labelled by ventral root injection of biocytin (green, C,E) were immuno-positive for the motoneuronal marker Islet 1-2 (white, B,D). F. Coronal slice of at the level of the mRN showing the injection site of TRDA on the right side. G-I. Coronal slices showing that TRDA injection (red) on the right mRN in F anterogradely labelled more fibers in the spinal cord ipsilaterally (I) than contralaterally (H). J. Confocal images showing that descending RS fibers (red) densely innervated the dendrites of motoneurons (green). K. Bar chart illustrating the optical density of TRDA immunofluorescence in the ventral spinal cord (spinal segment 3) ipsilateral versus contralateral to mRN injection measured in 10 spinal cord slices per animal from 3 animals. \*\*\**P* < 0.001, t test.

### Tectum evoke spinal motoneuron responses via the mRN

We then examined whether unilateral activation of the tectum translated into a contralateral spinal motor command. In an isolated brain-spinal cord preparation, we retrogradely labelled motoneurons on one side using Ca^2+^ green injections in the left ventral root of the third cervical segment. A stimulation electrode was placed in the tectum on the right side and the responses of spinal motoneurons were recorded (Fig. 6A-C). Trains of stimulation applied to the tectum evoked Ca^2+^ responses in motoneurons (n = 20 motoneurons pooled from 3 preparations, Fig. 6D-E). Increasing stimulation intensity increased the Ca^2+^ response amplitude in the motoneurons (Fig. 6D). Local application of glutamatergic antagonists in the mRN contralateral to the stimulated tectum decreased the spinal motoneuron responses (Fig. 6E). This indicated that tectum-evoked motoneuron activation are relayed by RS neurons in the mRN. Altogether, these experiments uncover a retino-tecto-reticulo-motoneuronal pathway through which retina activation is transformed into a motor command in salamanders. This pathway should play a key role in orienting behaviors in response to visual information (Fig. 7).

**Figure 6.**
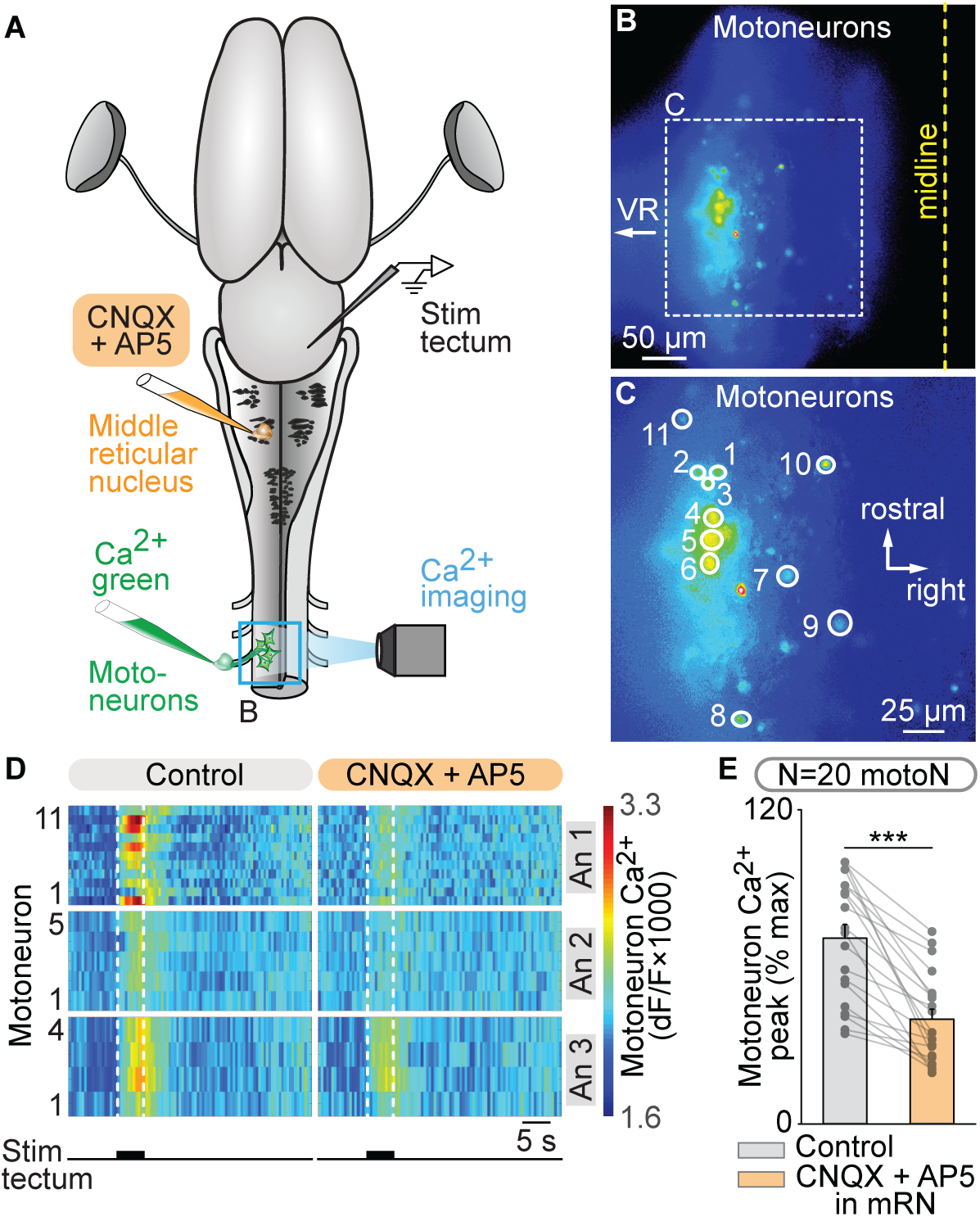
Tectum stimulation evokes spinal motoneuronal responses via a glutamatergic relay in the contralateral middle rhombencephalic nucleus (mRN). A. Schematic dorsal view of a salamander brain. A calcium (Ca^2+^) sensor (Ca^2+^ green) was applied onto the left ventral root of the third cervical spinal segment to retrogradely label motoneurons (motoN). A stimulation electrode was placed in the right tectum. The dorsal spinal cord was surgically removed on the left side to perform optical recordings of the motoneurons. B-C. Ca^2+^ fluorescence at rest of motoneurons. The spinal cord midline and the location of the ventral root (VR) are indicated. C is a magnification of the square in B. D. Color plots showing the Ca^2+^ responses of motoneurons (N = 20 motoN pooled from 3 animals, 4-15 cells per animal, An) to tectum stimulation (5 s train, 10 Hz, 2 ms pulses, 25-40 µA) before and 1-14 min after local injection of CNQX (1.9-5.7 pmol) and AP5 (1.0-2.9 pmol) in the mRN ipsilateral to the recorded motoneurons (and contralateral to the stimulated tectum) using a picospritzer (CNQX 1 mM, AP5 0.05 mM). Each line illustrates the average response of one cell (7 trials per cell per drug condition). Onset and offset of stimulation are indicated with vertical white dotted lines. Warm colors (red) indicate stronger Ca^2+^ responses. E. Bar charts showing the Ca^2+^ responses before and after bath application of the glutamatergic antagonists in the motoneurons. ***, *P* < 0.001 (paired t test).

**Figure 7.**
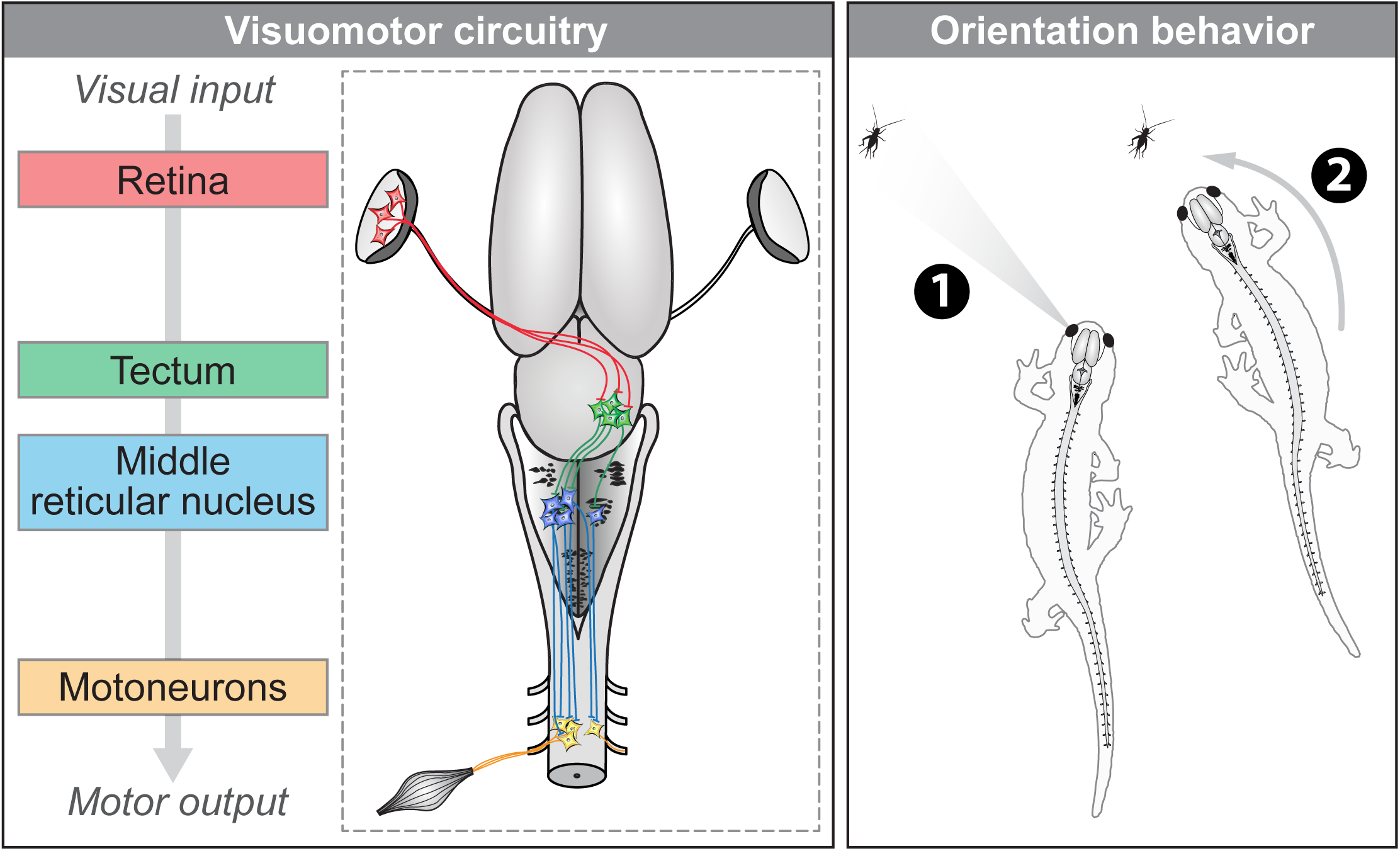
The proposed visuomotor circuitry in salamanders. **Left**, schematic dorsal view of a salamander brain showing the transformation of visual input into motor output. When retinal ganglion cells are activated in the left retina, tectal cells of the contralateral tectum are activated via a glutamatergic synapse. These tectal neurons send a stronger input to the reticulospinal (RS) neurons on the left side in the middle reticular nucleus via a glutamatergic synapse. From there, RS neurons send a mainly ipsilateral descending excitatory input to motoneurons, which increase muscular contraction on the side of the detected visual input, resulting in an orientation movement towards the visual stimulus. **Right**, the circuitry could account for prey tracking during ongoing locomotion. When a prey appears in the left visual field (e.g. cricket), the proposed circuitry would allow the animal to orient its body movements towards the prey and track it, to eventually snap it.

## Discussion

In the present study, we identified a neural substrate for visuomotor transformation in salamanders. The circuitry comprises the retina, tectum, mRN RS neurons and spinal motoneurons. Using tracing we show that retinal fibers project mainly to the contralateral tectum, where they contact the dendrites of tectal output neurons. These tectal neurons project to RS neurons of the mRN, a reticular nucleus known to control steering movements. mRN RS neurons were found to send mainly ipsilateral descending projections in apposition with the dendrites of cervical motoneurons. Using Ca^2+^ imaging we found that retina stimulation evoked stronger responses in ipsilateral mRN RS neurons. Microinjection of glutamatergic antagonists in the tectum contralateral to the stimulated retina decreased RS responses. Such tectal relay was confirmed in brain slices, where stimulation of the tectum superficial layer, where retinal inputs are located, evoked glutamatergic responses in deep tectal output neurons that projected to RS neurons. Tectum stimulation evoked stronger responses in contralateral mRN RS neurons. In a brainstem-spinal cord preparation, tectum stimulation evoked responses in contralateral cervical motoneurons, and injection of glutamatergic antagonists in the mRN contralateral to the stimulated tectum decreased motoneuron responses.

### A conserved visuomotor circuitry

The organization of the retino-tecto-reticulo-motoneuronal pathway we report here is in accordance with the circuitry proposed in salamander modeling studies to implement visually-guided orienting behaviors (Ijspeert and Arbib 2000, Petreska 2004). In our experiments, we do not provide a definitive evidence that the retino-tecto-reticulo-motoneuronal connections are monosynaptic. However, this is possible since in lamprey, retina send monosynaptic excitation to tectal output neurons (Kardamakis et al. 2015), tectal output neurons send monosynaptic input to RS neurons (Kardamakis et al. 2015), and RS neurons send monosynaptic input to motoneurons (Buchanan et al. 1987, Otah and Grillner 1989, Suzuki et al. 2019). We cannot rule out that spinal premotor interneurons could intervene in the motoneuronal responses. In lamprey, interneurons of the central pattern generator for locomotion are also targeted by RS neurons (Buchanan and Grillner 1987). In mice, spinal interneurons are targeted by ipsilaterally projecting Chx10^+^ RS neurons and this contributes to orienting maneuvers (Bouvier et al. 2015, Cregg et al. 2020, Usseglio et al. 2020). Therefore in salamanders RS connections on central pattern generator neurons could also contribute to tectum-evoked steering as proposed in a salamander modeling studies (Ijspeert and Arbib 2000, Petreska 2004) and also in a lamprey modeling study (Kozlov et al. 2014). Future studies should identify the muscles of the motoneurons recorded here, which possibly comprise motoneurons innervating axial or limb muscles (Francis 1934). Previous modeling studies suggested that targeting axial muscles is sufficient to evoke orienting manoeuvres during ongoing locomotion (Ijspeert and Arbib 2000). The descending RS projections possibly also innervate neck motoneurons, which are located in the cervical segments in salamanders (Nishikawa et al. 1991) as in birds (Hörster et al. 1990) and mammals (for review Isa and Sasaki 2002). Such possible connection to neck motoneurons would allow the animal to couple an ipsilateral head movement together with ipsilateral body bending towards the visual target, as recently shown in mice (Cregg et al. 2020, Usseglio et al. 2021).

It appears that the retino-tecto-reticulo-motoneuronal pathway is highly conserved in vertebrates. Lamprey and zebrafish tectal output neurons project to RS neurons, which send descending commands to spinal motor circuits (lamprey: Kardamakis et al. 2015, 2017, Suzuki et al. 2019; zebrafish: Gahtan et al. 2005). In mammals, descending connections from the superior colliculus (the mammalian tectum) to the RS system control turning and orienting movements in vivo (Cregg et al. 2020, Usseglio et al. 2021). These authors found that glutamatergic neurons of the superior colliculus project to contralateral Chx10^+^ RS neurons that control ipsilateral steering movements via ipsilateral projections to spinal cord neurons (Cregg et al. 2020, Usseglio et al. 2021). In our experiments, whether the RS neurons that responded to stimulation of the contralateral tectum also express Chx10 remains to be determined. This is possible since RS neurons projecting ipsilaterally are Chx10^+^ positive in mice (see e.g. Bouvier et al. 2015, Cregg et al. 2020, Usseglio et al. 2020) and the Chx10 gene is present in salamanders (Ryczko et al. 2020b).

Our study also highlights the multiple functions of RS neurons. In salamanders mRN RS neurons also receive the locomotor drive generated by the MLR, a region that controls locomotion initiation and speed in salamanders (Cabelguen et al. 2003, Ryczko et al. 2016a) as in other vertebrates (e.g. lamprey: Brocard et al. 2010, rat: Bachmann et al. 2013, cat: Shik et al. 1966, Opris et al. 2019; mouse: Bretzner and Brownstone 2013, Lee et al. 2014, Roseberry et al. 2016, Capelli et al. 2017, Caggiano et al. 2018, Josset et al. 2018, van der Zouwen et al. 2021, Dautan et al. 2021, pig: Chang et al. 2021). In lamprey, RS neurons also relay the MLR drive (Brocard et al. 2010), tactile inputs that evoke locomotion (Di Prisco et al. 1997), but also the olfactory inputs that evoke locomotion (Derjean et al. 2010, Daghfous et al. 2018, Beauséjour et al. 2020, for review Beauséjour et al. 2021). Altogether this strengthens the idea that RS neurons are key neurons that evoke and adapt locomotion in many circumstances and can be considered the “final common descending pathway” activating the spinal locomotor networks (Dubuc 2008).

### Orienting response vs. escape response

In the present study, retina stimulation evoked stronger ipsilateral responses at the RS level. Such activation pattern should evoke an orienting behavior towards the side of the stimulated retina. This is compatible with the ipsilateral body bending movements evoked by unilateral activation of mRN neurons in semi-intact preparations of salamanders (Ryczko et al. 2016c). Such asymmetric RS activity was shown to generate ipsilateral turning in a salamander robot controlled by a central pattern generator model driven by RS Ca^2+^ signals (Ryczko et al. 2016c). This is also consistent with the results of a mathematical model of the retino-tecto-reticulo-spinal pathway that controlled the movements of a simulated neuromechanical salamander tracking a prey (Ijspeert and Arbid 2000, Petreska 2004). In the present study, the stronger RS responses evoked by increased retina or tectum stimulations could be interpreted relative to the perceived position of the stimulus in the visual field (Petreska 2004). A modeling study suggested that more laterally located stimuli generate stronger orienting movements (Petreska 2004). How an increase in stimulation intensity in the retina or tectum translated into the recruitment of neurons differentially coding the position of the stimulus in the horizontal axis of the visual field remains to be determined. In lamprey, a stronger activation of the motor system ipsilaterally to the stimulated retina was also interpreted as an approach behavior (Kardamakis et al. 2015, Suzuki et al. 2019). A lamprey modeling study suggested that asymmetrical activity of tectal output neurons generate steering commands through modulation of RS activity during locomotion (Kozlov et al. 2014).

We were not able to evoke stronger mRN RS responses contralaterally to the stimulated retina, a response that would be interpreted as an escape response according to salamander modeling studies (Ijspeert and Arbid 2000, Petreska 2004) and previous lamprey studies (Kardamakis et all. 2015, Suzuki et al. 2019). A possibility is that our retina stimulations preferentially recruited tectal output neurons projecting to contralateral mRN RS neurons. In lamprey, such neurons are more excitable, with a spiking threshold 8 mV lower than those projecting ipsilaterally (Kardamakis et al. 2015). Future studies should determine whether a similar difference in intrinsic properties is present in salamander tectal output neurons. Another factor that could contribute is that we used electrical stimulation of the retina, which does not activate retinal cells the way a natural visual stimulus, such as a predator, would. In salamander retina slice, whole cell patch-clamp recordings showed that the brief current pulses used here (2 ms) preferentially activate retinal ganglion cells, which send their axons through the optic nerve to the brain, rather than local retinal bipolar or amacrine cells (Margalit and Thoreson 2006, Luo et al. 2012). The stimulation frequency used here (10 Hz) is in the appropriate range to activate retina ganglion cells (Kardamakis et al. 2015, for review Zueva 2018). In awake humans, electrical stimulation of the retina evokes perceived light spots (Humayun et al. 1999, Rizzo et al. 2003a and Rizzo et al. 2003b, Fan et al. 2019, for review Zueva 2018). Future studies should determine which patterns of light stimulation or natural images known to activate the salamander retina (Burkhardt et al. 2006, Burkhardt 2011) can evoke stronger response in ipsilateral RS neurons (orienting response) or in contralateral RS neurons (escape responses) (in lamprey see Kardamakis et al. 2015, Suzuki et al. 2019).

### Visuomotor processing beyond the optic tectum

On top of the retino-tecto-reticulo-motoneuronal pathway we describe here, other circuits likely contribute to visuomotor processing in salamanders. In the tectum, local GABA-immunoreactive neurons are present in all the cellular layers (Franzoni and Morino 1989). In lamprey, tectal GABAergic interneurons play a role in the selectivity of the activated tectal output neurons (Kardamakis et al. 2015). Beyond visual information, the tectum receives other sensory modalities and performs multisensory integration, such as aligning auditory and visual maps in owls (e.g. Knudsen 1982, Reches and Gutfreund 2009, Mysore et al. 2010, modeled in Huo et al. 2012) or integrate visual and electrosensory information in lamprey (Kardamakis et al. 2016). Future studies should explore whether the pathways reported here can be controlled by GABAergic tectal transmission and integrate multisensory information.

The salamander tectum is innervated by neuromodulatory systems as shown by the presence of e.g. serotoninergic fibers (Gruber and Harris 1981, Dicke et al. 1997), cholinergic fibers (Marin and Gonzalez 1999, Sanchez-Camacho et al. 2002) and dopaminergic fibers (Sanchez-Camacho et al. 2002). These inputs likely influence visuomotor processing in salamanders. In other species, if we consider dopaminergic inputs alone for instance, dopaminergic fibers originating from the posterior tuberculum increase the excitability of tectal output neurons via D1 receptor activation and decrease it via D2 receptor activation (Perez-Fernandez et al. 2017, von Twickel et al. 2019). In toads the dopaminergic agonist apomorphine attenuates prey-oriented turning movements (Glagow and Ewert 1997). In mammals, dopaminergic fibers from A13 innervate the superior colliculus and likely influence the ability to allocate attention and orienting movements towards regions of the visual field (Bolton et al. 2015, Woolrych et al. 2021).

The pretectum might play a crucial role in the switch between orienting or avoidance responses. This idea comes from dual lesion paradigms in toads. After a first lesion (bilateral ablation of the telencephalon, i.e. striatum and pallium), no visually-guided prey catching occurs (Matsumoto et al. 1991). Strikingly, adding a second lesion in the pretectum results in “disinhibition” of orienting behaviors, i.e. increased prey catching towards any moving object. Such effect of pretectum lesion was also observed in salamanders (Finkenstädt 1980). Future studies should study whether and how the pretectum controls the pathway we here report.

The innervation of the thalamus that we report likely underlies the previously reported responses evoked by optic nerve stimulation in thalamic neurons, that relay visual information higher brain regions in salamanders (Roth and Grunwald 2000). Such retino-thalamo-pallial pathway, well known in mammals (Butler 1994a) birds (Butler and Hodos, 1996) and frogs (Neary and Northcutt 1983), was also recently described in the phylogenetically older lamprey (Suryanarayana et al. 2020) and is therefore another conserved component of the visual system. It was proposed to play a role in sensory representation and cognition-based visuomotor processing in lamprey (Suryanarayana et al. 2020). In salamanders, the thalamus receives projections from the optic nerve and relay the information to the pallium, septum, striatum and amygdala (Ruhl and Dicke 2012). The thalamus is also bilaterally connected with the tectum (Ruhl and Dicke 2012). Lesion of the dorsal thalamus in salamanders impairs recognition, evaluation of objects and spatial attention. Salamanders can still snap a prey presented in front of their head, but in the presence of two prey objects, they are less able to select one and orient towards it (Rulh and Dicke 2012). Future studies should examine how these higher centers control the pathway reported here.

### Conclusion

We uncovered a retino-tecto-reticulo-motoneuronal circuitry through which visual input can be transformed into a spinal motor command in salamanders. This pathway is likely involved in orienting behaviors during prey capture. Our work shows that the organisation of visuomotor pathways is largely similar to that of lamprey, zebrafish and mammals. Intriguingly, the salamander has the extraordinary capacity to regenerate its nervous system (for review Joven and Simon 2018). The retina of salamanders, which has been extensively studied at the cellular level (for review Thoreson 2021), regenerates and regrows its central projections after lesion (Okamoto et al. 2007). Thus, salamanders provide a unique opportunity to study how visuomotor pathways are repaired after lesion in tetrapods.

## Acknowledgments

We thank Jean Lainé for technical assistance with the microscopy platform, Alberto Joven and Andras Simon for the animals and Jordan Swiegers for the help with some of the experiments.

## Author Contributions

AF: conceptualization, investigation, data curation, formal analysis, methodology, validation, visualization, writing—review and editing. DR: conceptualization, investigation, data curation, formal analysis, funding acquisition, methodology, project administration, resources, supervision, validation, visualization, writing—original draft, writing—review and editing. All authors contributed to the article and approved the submitted version.

## Data availability

All relevant data are available from the authors.

